# PredHLM: quantitative and interpretable prediction of metabolic half-life in human liver microsomes

**DOI:** 10.64898/2026.07.02.736062

**Authors:** Jidon Jang, Nam-Chul Cho, Jae Yong Ryu, Kwang-Seok Oh

**Affiliations:** Data Convergence Drug Research Center, Korea Research Institute of Chemical Technology, 141 Gajeong-ro, Yuseong-gu, Daejeon 34114, Republic of Korea; Drug Information Platform Center, Korea Research Institute of Chemical Technology, 141 Gajeong-ro, Yuseong-gu, Daejeon 34114, Republic of Korea; Department of Medicinal and Pharmaceutical Chemistry, University of Science and Technology, 217 Gajeong-ro, Yuseong-gu, Daejeon 34113, Republic of Korea; School of Systems Biomedical Science, Soongsil University, 369 Sangdo-Ro, Dongjak-Gu, Seoul 06978, Republic of Korea

**Keywords:** Metabolic stability, Machine learning, Human liver microsome, Half-life prediction, Interpretability, Drug discovery

## Abstract

Human liver microsome (HLM)-based metabolic stability assays are fundamental in early drug discovery, shaping pharmacokinetic profiles and oral bioavailability. However, these experimental assays are labor-intensive and time-consuming, limiting their application in large-scale virtual screening. Computational models can prioritize compounds at scale, yet most are classification-based, leaving quantitative and interpretable prediction of HLM half-life limited. In this study, we developed a quantitative machine learning model for the direct prediction of HLM half-life (T_1/2_) by integrating 11,790 compounds combining in-house and curated public data. Among various combinations of molecular features and learning algorithms, the XGBoost model with RDKit 2D descriptors achieved the best predictive performance, with an RMSE of 0.507 and an R^2^ of 0.431 on an independent test set. Shapley Additive Explanations (SHAP) analysis identified lipophilicity and known metabolic soft-spot features as the primary contributors to the predictions. These results suggest that this quantitative approach provides a practical framework for defining metabolic stability margins, thereby supporting rapid Go/No-go decisions in preclinical drug discovery. The source code, data, and trained model are available at https://github.com/joshua-416/PredHLM.

**Scientific Contribution:** This study integrates in-house and curated public data for 11,790 compounds to enable quantitative HLM half-life prediction beyond classification-based metabolic stability assessment. Comparative evaluation of molecular representations and learning algorithms identifies XGBoost with RDKit 2D descriptors as the best-performing configuration among those tested. SHAP analysis links predictions to lipophilicity and known metabolic soft-spot features, while an applicability domain based on the sum of distance-weighted contributions (SDC) supports compound-specific assessment of prediction reliability and compound prioritization.

**Graphical Abstract:** 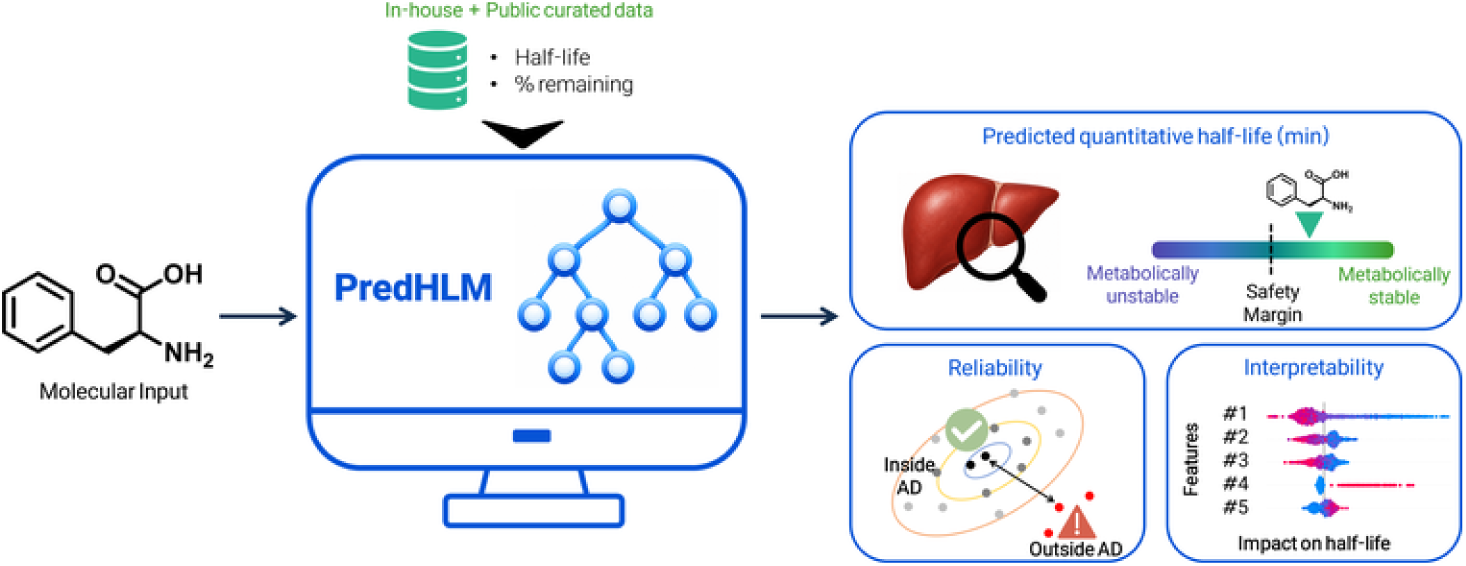

## Introduction

Metabolic stability is a critical pharmacokinetic determinant that governs systemic exposure, clearance rate, half-life, and oral bioavailability. In the early stages of drug discovery, insufficient metabolic stability often leads to high attrition rates or an increased burden during lead optimization. Human liver microsomes (HLM), containing various cytochrome P450 enzymes, are the gold-standard in vitro platform for evaluating phase I metabolism [1]. Specifically, the metabolic half-life derived from HLM assays provides an intuitive, concentration-normalized parameter for estimating the metabolic rate. However, experimental HLM assays are labor-intensive and resource-heavy, making them a bottleneck for the rapid evaluation of large chemical libraries.

To reduce this experimental burden, various quantitative structure-activity relationship (QSAR) and machine learning (ML) models have been developed. However, several critical gaps remain in HLM stability prediction. First, the majority of existing models are restricted to classification tasks (e.g., stable vs. unstable), which lack the granularity required for precise lead optimization [2–9]. Second, quantitative half-life measurements are subject to a non-negligible experimental uncertainty. Even in a standardized single-laboratory microsomal stability assay, control compounds showed a minimum significant ratio of approximately two-fold [10]. This uncertainty is further amplified in public datasets by heterogeneous protocols, inter-laboratory variability, and censoring at the stability extremes [11]. This noise imposes a practical ceiling on regression accuracy and has led many researchers to adopt classification, as regression performance deteriorates markedly under rigorous time-split or external validation [12]. Third, many black-box models lack interpretability, failing to provide chemical insights that align with experimentally validated metabolic soft-spots.

Within this landscape, prior quantitative regression models have generally targeted related endpoints such as human hepatic clearance or percent remaining at a single time point [12, 13], or been confined to narrow chemotypes such as arylpiperazine derivatives [14]. Even when microsomal half-life is the nominal endpoint, it is often dichotomized into stable/unstable categories rather than predicted as a continuous value [15, 16]. More recently, MetaStab-Analyzer introduced multi-species microsomal half-life regression on publicly available data, with modest reported accuracy for HLM (R^2^ ≈ 0.40) [17]. Direct and interpretable prediction of continuous HLM half-life, coupled with reliability estimation, therefore remains underexplored, despite its value for discriminating close analogs and defining safety margins.

In this study, we present a quantitative regression framework for predicting HLM half-life, leveraging a curated dataset of 11,790 compounds. This dataset integrates in-house Korea Chemical Bank (KCB) data with curated public data. We benchmarked multiple machine learning and deep learning architectures. To overcome the ‘black-box’ nature of ML, we employed Shapley Additive Explanations (SHAP) [18] to elucidate the molecular features driving the prediction, ensuring alignment with known metabolic pathways. We also assessed prediction reliability using an applicability domain metric. Finally, we release the source code, curated datasets, and trained model on GitHub to support practical application and reproducibility.

## Methods

### Data collection

Experimental metabolic stability data for HLM were integrated from both in-house and public repositories. The in-house dataset was curated from the Korea Chemical Bank (KCB), comprising 3,360 records of experimentally measured half-life (T_1/2_) and 3,954 records of percent remaining at 30 min, measured by LC-MS/MS at an initial 2 μM substrate concentration. To expand the chemical space, public data were retrieved from ChEMBL (v36), PubChem (AIDs: 1555, 624452, 624142, and 1963597).

To ensure consistency across diverse sources, percent remaining data at T min (R_T_) were converted into half-life (T_1/2_) values based on a first-order kinetic equation:

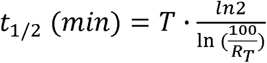

where T represents the incubation time (min), and *R_T_* is the percentage of the parent compound remaining at T min.

### Data curation and quality control

Following data collection and endpoint harmonization, a multi-step curation procedure was applied to minimize assay heterogeneity and improve measurement reliability. For ChEMBL records, only assays consistent with HLM-based parent compound depletion or half-life measurements were retained. Records were excluded if they corresponded to non-HLM systems, including hepatocytes, cytosol, S9 fraction, plasma, and intestinal assays, or represented assays and conditions outside phase I HLM depletion (full criteria in Supplementary Methods). Records with missing numerical values, unsupported units, or insufficient assay-description evidence were also removed. PubChem BioAssay records were manually inspected to retain only HLM metabolic stability assays not originating from ChEMBL.

For percent remaining data, records reported as turnover, transformation, disappearance, or compound metabolized were interpreted as parent compound depletion and converted to percent remaining by subtracting the reported value from 100. Inequality signs were reversed accordingly. Values reported as 0 or ≥100 were replaced with 0.001 and 99.999, respectively, to enable numerical conversion. Censored records were retained only when they provided clear stability information: percent remaining bounds in extreme ranges, such as <10% or >90%, and half-life bounds sufficiently distant from the 30 min stability threshold were used, whereas ambiguous censored records were excluded.

After integration of the curated half-life and converted stability records, duplicated compounds were represented by the arithmetic mean of their half-life values. This process yielded 12,467 unique compounds before outlier filtering. Half-life values were then log10-transformed to reduce their strong right skew and stabilize variance, and extreme upper outliers were removed using the interquartile range method with k = 1.5 to improve training stability. The final dataset used for model development contained 11,790 compounds.

The distribution of curated HLM half-life values is shown in Fig. 1a. Structural diversity was assessed by t-SNE of ECFP4 fingerprints (Fig. 1b): public datasets tended to form localized clusters, whereas the in-house KCB compounds provided broader and more uniform coverage of the chemical space.

**Figure 1.**
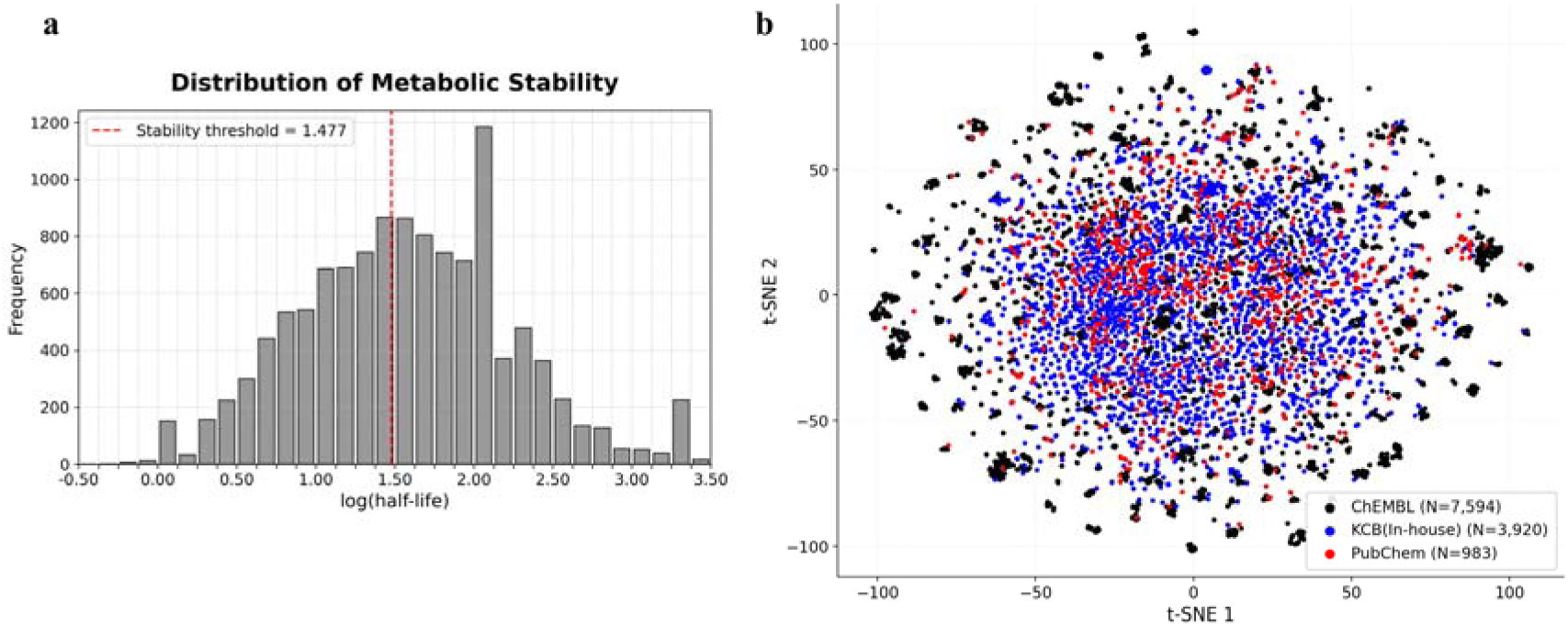
Distribution and chemical-space coverage of the curated HLM half-life dataset. **a** Histogram of log10-transformed HLM half-life values after data curation. The red dashed line denotes the 30 min half-life stability threshold, a cutoff widely used to label compounds as metabolically stable or unstable in previous studies [8, 11]. **b** t-SNE visualization based on 2,048-bit ECFP4 fingerprints with radius 2, showing the chemical-space distribution of in-house and public compounds.

### Molecular featurization and model architectures

The overall model development workflow is illustrated in Fig. 2. Four featurization schemes were evaluated: ECFP4 fingerprints (2,048-bit) [19], RDKit 2D descriptors (http://www.rdkit.org), Mordred descriptors [20], and MACCS keys [21]. Descriptors with missing or non-finite values were excluded, and the RDKit Ipc descriptor was removed because it produced extreme numerical values for a subset of molecules. The retained RDKit 2D and Mordred descriptors are listed in Table S1. These representations were utilized to benchmark nine predictive architectures, including multilayer perceptron (MLP) [22], k-nearest neighbors (k-NN) [23], kernel ridge regression (KRR) [24], support vector machines (SVM) [25], and tree-based algorithms such as random forest (RF) [26], LightGBM [27], XGBoost [28], and CatBoost [29]. Additionally, a communicative message passing neural network (CMPNN) [30] was employed to capture intrinsic structural features directly from molecular graphs. Building on this base CMPNN, we evaluated three variants: a self-attention [31] readout applied after message passing, concatenation of precomputed molecular features with the graph embedding before a fully connected prediction layer, and a combination of both. The feature-augmented variants were evaluated across all four featurization schemes.

**Figure 2.**
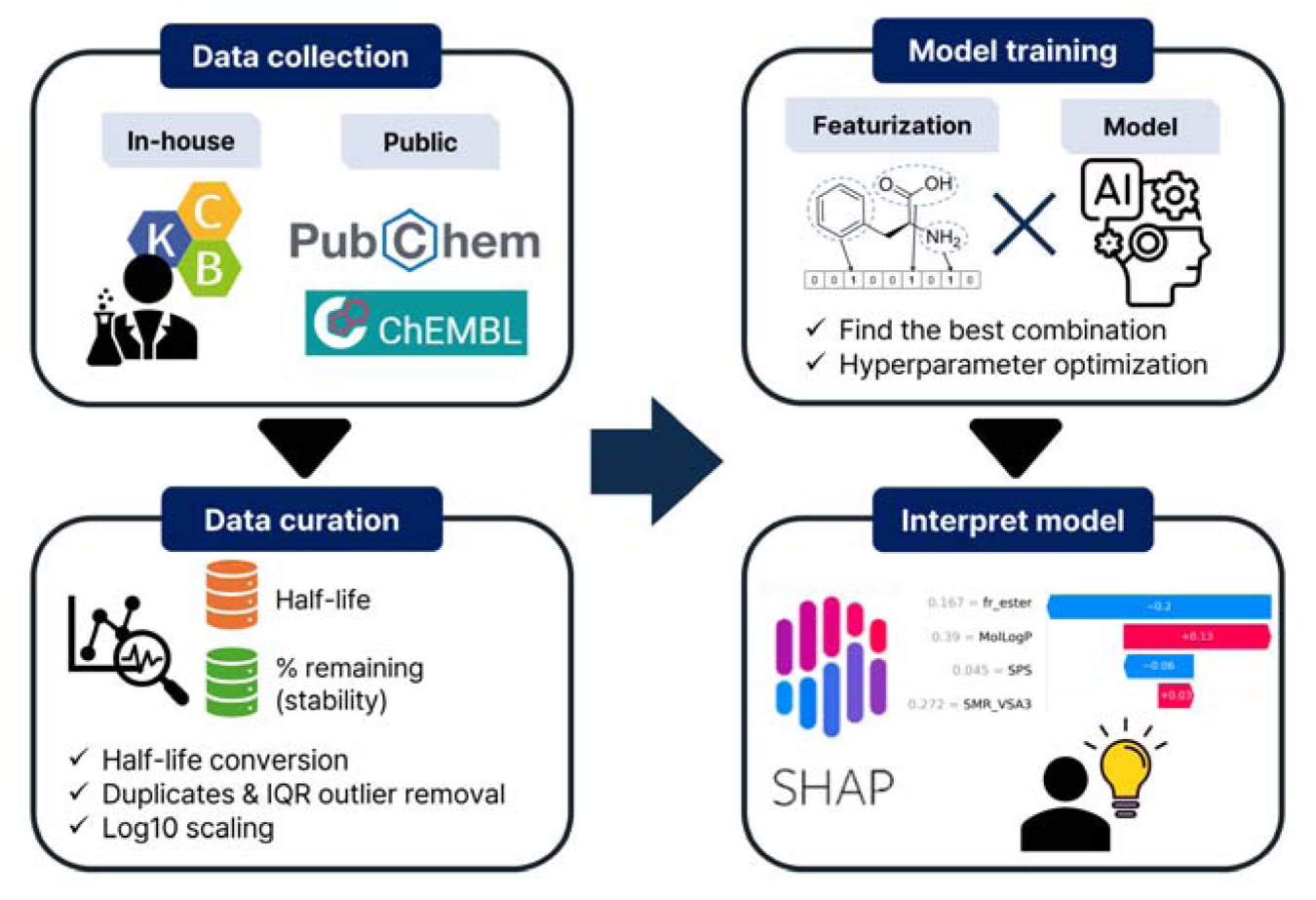
Schematic diagram for overall model development process.

### Training and evaluation protocol

The curated dataset was partitioned into a training set and an independent test set at a ratio of 9:1 using a random split. All hyperparameter optimization and model selection were performed within the training set using 5-fold cross-validation, whereas the test set was reserved exclusively for the final evaluation of the selected model. For continuous descriptor-based feature sets, including RDKit 2D and Mordred descriptors, features were standardized using a StandardScaler fitted on the training folds only, and the same transformation was applied to the held-out fold and the independent test set to prevent data leakage. For each feature-algorithm combination, hyperparameters were optimized using Optuna [32] for 100 trials, with the mean cross-validated mean squared error (MSE) across the five folds used as the objective function. For CMPNN and its variants, whose greater per-trial computational cost limited the feasible search budget, 40 trials were used instead. The hyperparameter search spaces are summarized in Table S2, and the final hyperparameters selected for each model are reported in Table S3 and S4. The best-performing feature-algorithm combination was selected as the final model. For this configuration, the five cross-validation fold models were aggregated into an ensemble whose averaged predictions were evaluated on the independent test set. Predictive performance was assessed using root mean squared error (RMSE), mean absolute error (MAE), and the coefficient of determination (R^2^). For the selected ensemble, Shapley Additive Explanations (SHAP) analysis (Lundberg and Lee, 2017) was performed to quantify feature-level contributions.

Beyond this random-split evaluation, the selected model was additionally evaluated under a scaffold-based split to assess generalization to structurally novel chemotypes. Following the MoleculeNet benchmarking strategy [33], molecules were grouped by their Bemis–Murcko frameworks [34] computed with RDKit, and whole scaffold groups were assigned exclusively to either the training or the test set so that no scaffold was shared between them, with groups allocated until the test set comprised 10% of the data under a fixed random seed. The inner 5-fold cross-validation used for hyperparameter optimization and ensemble construction was likewise performed at the scaffold level using GroupKFold from scikit-learn [35], ensuring that no scaffold was split across folds. Because no test compound shares a Bemis–Murcko framework with any training compound, this protocol provides a conservative, leakage-free estimate of prospective predictive performance.

### Metric for applicability domain

To assess prediction reliability, the applicability domain (AD) was evaluated using the sum of distance-weighted contribution (SDC) approach. The SDC index quantifies how well a query compound is represented within the training chemical space. For this analysis, molecules were encoded using 2,048-bit ECFP4 fingerprints, and the structural similarity between a query compound and each molecule in the training set was quantified using the Tanimoto coefficient [36]. The SDC score for a query compound was defined as follows [37]:

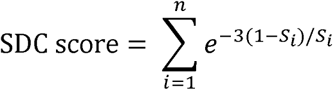

where *S_i_* denotes the Tanimoto similarity between the query compound and the *i*-th compound in the training set, and *n* represents the total number of compounds in the training set. Because contributions are distance-weighted, molecules in densely populated regions of the training chemical space receive higher SDC scores, whereas structurally distant compounds receive lower scores. Thus, lower SDC values flag compounds outside the applicability domain, whose predictions are less reliable.

## Results and Discussion

### Model benchmarking and performance metrics

Eight machine learning algorithms were evaluated across four featurization schemes using 5-fold cross-validation on the training set. Among all tested combinations, XGBoost with RDKit 2D descriptors achieved the best performance, with a mean cross-validated RMSE of 0.515 and R^2^ of 0.410. The performance of models using RDKit 2D descriptors is summarized in Table 1. Full benchmarking results for the remaining three featurization schemes (Mordred descriptors, ECFP4 fingerprints, and MACCS keys) are provided in Table S5. Although the Mordred set attained a marginally lower cross-validated RMSE (by 0.001, within the cross-validation standard deviation and thus not meaningful), RDKit 2D descriptors were retained for parsimony and interpretability: the set is far more compact (∼800 fewer descriptors), reducing computational cost and overfitting risk, while its physically intuitive descriptors map more readily onto medicinal-chemistry concepts and support the SHAP-based interpretation central to this study.

**Table 1.** Cross-validated regression performance of models using RDKit 2D descriptors for HLM half-life prediction.

| Algorithm | RMSE (↓) | MAE (↓) | R<sup>2</sup> (↑) |
| --- | --- | --- | --- |
| MLP | 0.557 ± 0.009 | 0.422 ± 0.007 | 0.311 ± 0.022 |
| k-NN | 0.550 ± 0.005 | 0.415 ± 0.007 | 0.328 ± 0.013 |
| KRR | 0.538 ± 0.008 | 0.406 ± 0.007 | 0.357 ± 0.017 |
| SVM | 0.526 ± 0.005 | 0.392 ± 0.005 | 0.384 ± 0.009 |
| RF | 0.529 ± 0.005 | 0.402 ± 0.006 | 0.377 ± 0.008 |
| LightGBM | 0.519 ± 0.007 | 0.389 ± 0.006 | 0.401 ± 0.012 |
| CatBoost | 0.517 ± 0.006 | 0.387 ± 0.006 | 0.406 ± 0.009 |
| **XGBoost** | **0.515 ± 0.007** | **0.387 ± 0.007** | **0.410 ± 0.011** |
| CMPNN | 0.527 ± 0.003 | 0.397 ± 0.003 | 0.382 ± 0.010 |
| CMPNN + attention | 0.534 ± 0.003 | 0.404 ± 0.006 | 0.367 ± 0.009 |
| CMPNN + RDKit 2D | 0.526 ± 0.003 | 0.395 ± 0.004 | 0.386 ± 0.013 |
| CMPNN + attention + RDKit 2D | 0.528 ± 0.004 | 0.400 ± 0.004 | 0.380 ± 0.014 |

Values are reported as mean ± standard deviation across 5-fold cross-validation; the best-performing model is shown in bold

We further evaluated four CMPNN-based variants to assess the contribution of graph-based representation learning. Among these models, CMPNN with RDKit 2D descriptors showed the best graph-based performance, with an RMSE of 0.526 and R^2^ of 0.386, but it did not outperform the optimized XGBoost model. Adding RDKit 2D descriptors to CMPNN slightly improved performance, whereas adding attention did not. These results suggest that, for the present HLM dataset, global physicochemical and topological descriptors captured the relevant structure-stability relationships more effectively than graph-based representation learning.

The cross-validation ensemble of the selected RDKit 2D-XGBoost configuration was subsequently applied to the independent test set, achieving an RMSE of 0.507 and an R^2^ of 0.431 (Fig. 3). As shown in the parity plot, the predictions captured the overall trend of the experimental values, although the model showed a slight tendency to overestimate compounds with very short half-lives and to underestimate highly stable compounds. This trend is likely related to the lower sample density at both extremes of the half-life distribution (Fig. 1a), which limits model calibration in these regions. Nevertheless, the optimized model compared favorably with a previous half-life model (RMSE of 0.64, R^2^ of 0.40) [17].

**Figure 3.**
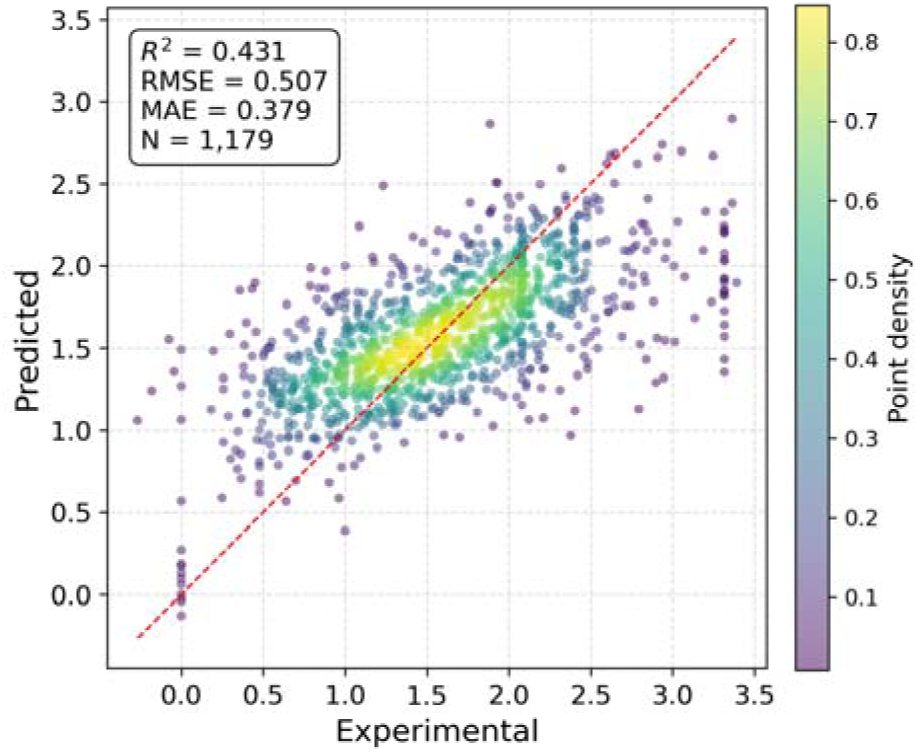
Parity plot of experimental versus predicted log-transformed HLM half-lives for the optimized RDKit 2D-XGBoost ensemble on the independent test set. Point color indicates local point density estimated by Gaussian kernel density estimation, highlighting regions where test compounds are densely distributed. The red dashed line denotes the identity line (y = x).

To further probe generalization to structurally novel chemotypes, the selected RDKit 2D-XGBoost model was re-evaluated under the scaffold-based split. Performance decreased only modestly (cross-validated RMSE of 0.545 ± 0.005; independent test RMSE of 0.536; full metrics in Table S6), consistent with generalization to structurally novel chemotypes rather than reliance on analog-specific memorization.

### Characterization of the prediction reliability via the applicability domain

The applicability domain of the optimized XGBoost model was evaluated on the independent test set using the SDC score. As shown in Fig. 4a, the residuals of the full test set (n = 1,179) were generally centered around zero, but compounds with low SDC values showed broader error distributions and more frequent large deviations, whereas high-SDC compounds were more tightly distributed around zero.

**Figure 4.**
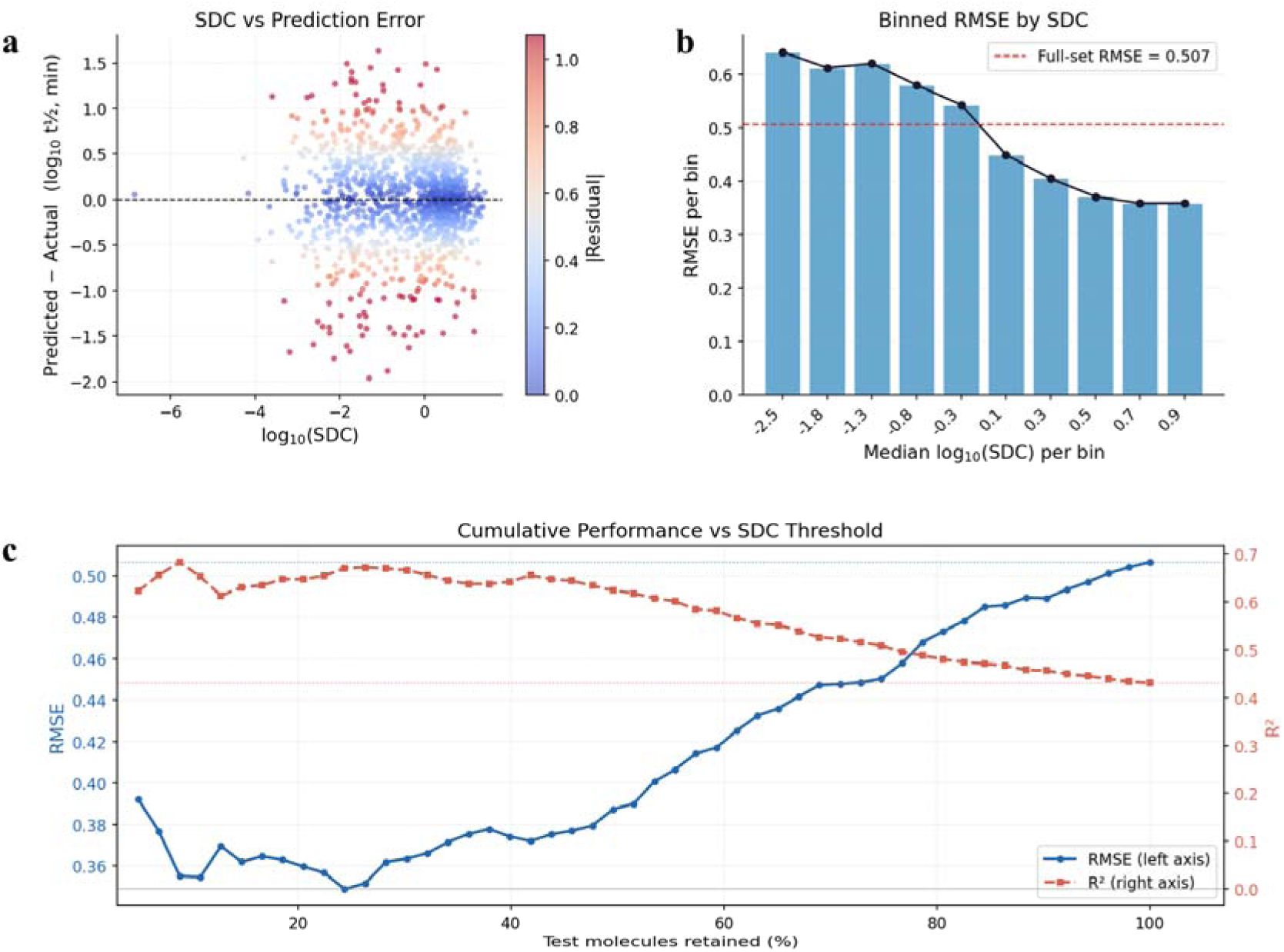
Applicability domain analysis based on the SDC score. **a** Relationship between log_10_(SDC) and prediction residuals on the independent test set. Residuals were calculated as predicted minus experimental log_10_ half-life, and point color indicates the absolute residual. **b** RMSE across ten equal-frequency SDC bins. The dashed red line indicates the full test-set RMSE. **c** Cumulative RMSE and R^2^ after ranking test compounds by decreasing SDC.

Consistent with this trend, the binned error analysis in Fig. 4b showed that RMSE decreased as the median log_10_(SDC) increased. Low-SDC bins exhibited RMSE values above the full-set baseline, while high-SDC bins showed lower prediction errors. These results indicate that compounds located in well-represented regions of the training chemical space were predicted more accurately.

The cumulative performance analysis in Fig. 4c further supports the use of SDC as a prediction confidence indicator. When test compounds were ranked by decreasing SDC, the highest-confidence subset showed improved performance, reaching a maximum R^2^ of 0.683 for the top 8.9% of compounds. As lower-SDC compounds were progressively included, RMSE increased and R^2^ approached the full-set values. Overall, the SDC-based applicability domain reflects prediction reliability and can flag compounds whose predictions warrant greater caution.

### External benchmarking against classification models

We next benchmarked the optimized model against published models on an external test set compiled from two datasets reported by Ryu et al. and Aliagas et al. [7, 12]. Because the available reference tools output binary stability classes rather than continuous half-lives, PredHLM’s continuous predictions were correspondingly binarized for direct comparison [7, 11]. Compounds were defined as metabolically stable when the HLM half-life exceeded 30 min, corresponding to more than 50% parent compound remaining after 30 min under first-order disappearance kinetics.

To prevent data leakage, compounds overlapping with the PredHLM training set were removed from the external test set, yielding a shared evaluation set of 958 compounds on which all models were assessed. The predicted half-life values were binarized at the 30 min cutoff for threshold-dependent metrics, including accuracy, positive/negative predictive value (PPV/NPV), sensitivity, specificity, F1-score, and Matthews correlation coefficient (MCC). ROC-AUC was calculated directly from the continuous predicted half-life values as ranking scores against the binary experimental labels, without applying a fixed threshold. For the reference models, predictions were generated on the same 958-compound evaluation set using their publicly available web-tools. Because the exact training compounds of these models were not disclosed, any residual overlap with their training data could not be excluded and would, if present, favor the reference models.

As summarized in Table 2, PredHLM achieved the highest accuracy, PPV, NPV, sensitivity, MCC, F1-score, and ROC-AUC among the compared models. Although its specificity was slightly lower than that of the best-performing reference models, the improved MCC and F1-score indicate more balanced classification performance. The higher sensitivity and NPV indicate improved detection of metabolically stable compounds.

**Table 2.**
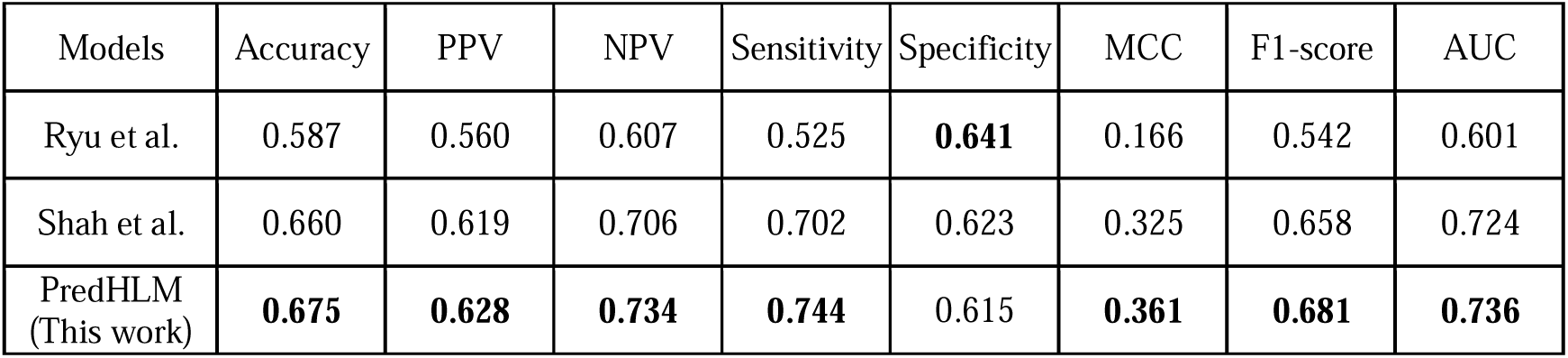
External classification performance of PredHLM and reference HLM metabolic stability prediction models.

| Models | Accuracy | PPV | NPV | Sensitivity | Specificity | MCC | F1-score | AUC |
| --- | --- | --- | --- | --- | --- | --- | --- | --- |
| Ryu et al. | 0.587 | 0.560 | 0.607 | 0.525 | **0.641** | 0.166 | 0.542 | 0.601 |
| Shah et al. | 0.660 | 0.619 | 0.706 | 0.702 | 0.623 | 0.325 | 0.658 | 0.724 |
| PredHLM<br>(This work) | **0.675** | **0.628** | **0.734** | **0.744** | 0.615 | **0.361** | **0.681** | **0.736** |

Unlike conventional classification models, PredHLM directly predicts quantitative HLM half-life values. This regression-based output preserves continuous stability information, allows flexible application of alternative decision thresholds, and provides a more interpretable estimate of metabolic stability margin. Thus, PredHLM can support both binary Go/No-go decisions and finer compound prioritization during lead optimization.

### SHAP-based interpretation of the optimized model

To elucidate the decision-making process of the optimized XGBoost model, SHAP analysis was applied to the independent test set predictions. Positive and negative SHAP values indicate a corresponding increase or decrease in the predicted log-transformed half-life relative to the baseline. Global feature importance and per-compound feature effects are shown in Fig. 5a and 5b, respectively.

**Figure 5.**
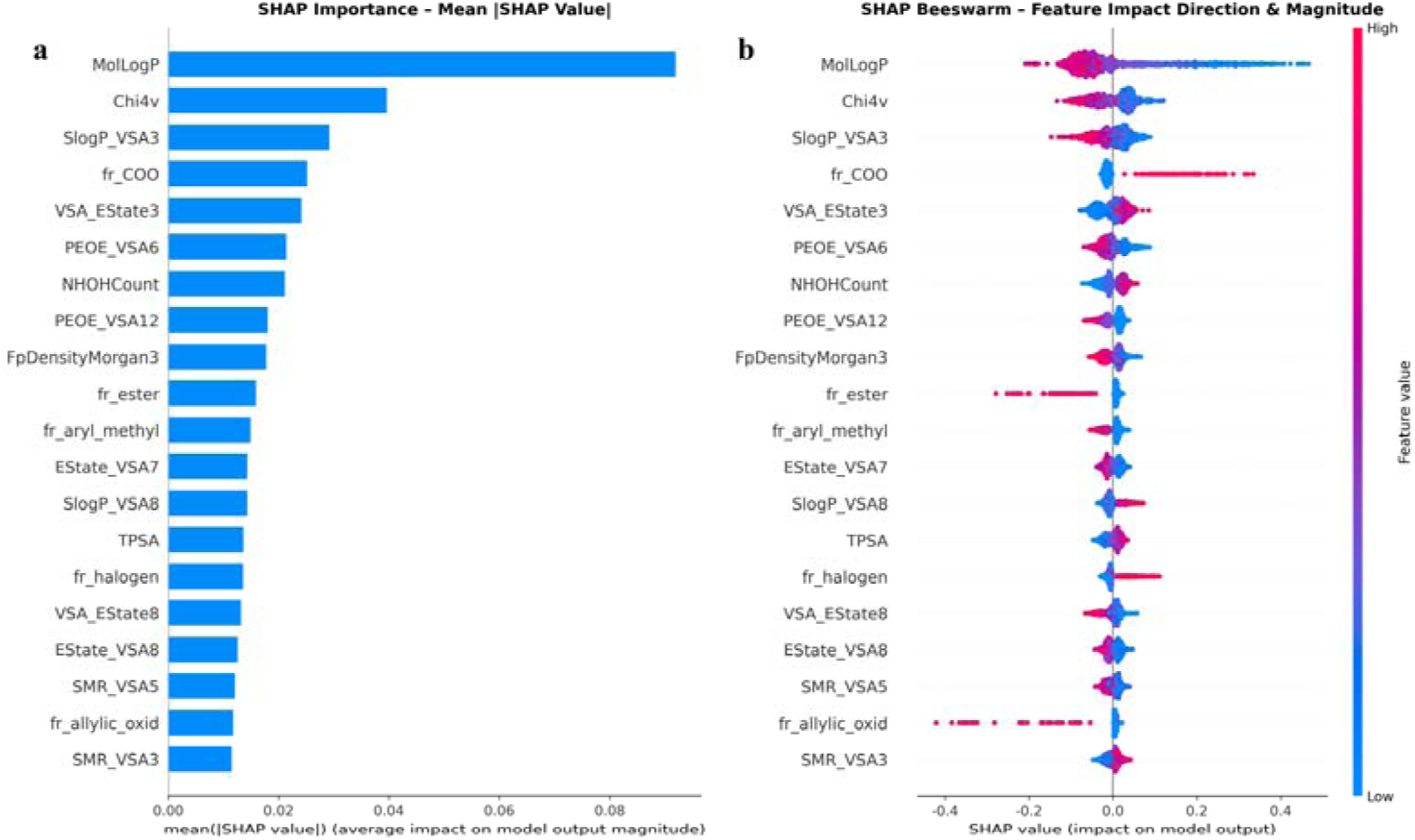
SHAP analysis of the optimized RDKit 2D-XGBoost model on the independent test set. **a** Global feature importance of the top 20 descriptors ranked by mean absolute SHAP value. **b** Beeswarm summary plot showing the distribution and direction of feature-level SHAP contributions across individual test-set compounds, with point color encoding the descriptor value (red, high; blue, low).

MolLogP emerged as the most influential descriptor, identifying lipophilicity as a primary determinant of HLM stability. Higher lipophilicity is associated with shorter predicted half-lives (negative SHAP values). This aligns with the established pharmacokinetic principle that increased lipophilicity enhances microsomal partitioning and susceptibility to CYP-mediated oxidative metabolism [38, 39]. Conversely, descriptors related to polarity and hydrogen-bonding capacity, such as TPSA and NHOHCount, exhibited positive associations with half-life, suggesting that these features may reduce susceptibility to rapid phase I microsomal metabolism in this dataset.

The model also identified several metabolic soft-spots as critical fragment descriptors. The aryl methyl group (fr_aryl_methyl) and allylic oxidation sites (fr_allylic_oxid), known for their susceptibility to CYP-mediated oxidation [40, 41], showed a negative contribution to predicted half-life. Similarly, the ester group descriptor (fr_ester) was associated with decreased stability, reflecting its vulnerability to hydrolysis by carboxylesterases [42]. In contrast, the halogen count descriptor (fr_halogen) showed a positive association with half-life, consistent with the established strategy of using halogenation, particularly fluorination, to block metabolic soft spots and improve stability [43].

Interestingly, the carboxylic acid group (fr_COO) was positively associated with half-life. While carboxylic acids are subjected to UGT-mediated acyl glucuronidation, they often exhibit lower susceptibility to CYP-mediated oxidative pathways due to their high polarity and ionization under physiological conditions [44]. Given that our dataset was curated to focus on phase I HLM assays where phase II co-factors (e.g., UDPGA) were excluded, the model correctly identified carboxylic acids as relatively stable features in this context. Other highly ranked descriptors, including Chi4v, FpDensityMorgan, and several binned VSA-type descriptors (PEOE_VSA*, EState_VSA*, VSA_EState*, SMR_VSA*, and SlogP_VSA*, where the asterisk denotes multiple members of each family), encode broader molecular properties such as topology, structural complexity, and surface-area-weighted physicochemical characteristics. Although these descriptors also contributed to model predictions, their direct mechanistic interpretation is less straightforward and should therefore be considered supportive rather than causal.

Overall, the SHAP analysis suggests that the model captures chemically meaningful structure-stability relationships. The alignment between model-derived feature importance and established medicinal chemistry principles, including lipophilicity and specific metabolic soft-spots, supports the interpretability of the model.

## Conclusion

In this study, we developed PredHLM, an interpretable machine learning framework for the quantitative prediction of HLM metabolic half-life. By integrating in-house experimental data with curated public datasets, the model provides direct half-life estimates rather than only categorical stable/unstable labels, enabling more granular compound prioritization during lead optimization. The optimized descriptor-based model achieved competitive predictive performance and generalized to structurally novel chemotypes under a stringent scaffold-based split. In external validation, its regression output also supported reliable classification. Applicability domain analysis further provided a practical confidence measure, and SHAP interpretation suggested that the model captured chemically plausible determinants of HLM stability such as lipophilicity and metabolic soft-spot fragments.

Nevertheless, PredHLM should be regarded as a decision-support tool rather than a replacement for experimental HLM assays. The model remains affected by assay variability, censored measurements, and uncertainty introduced by converting percent remaining values into half-life under first-order kinetic assumptions. In addition, the current dataset primarily reflects phase I microsomal metabolism and does not fully represent phase II metabolism, hepatocyte-specific pathways, transporter effects, or in vivo pharmacokinetics. Future work will focus on prospective experimental validation, expansion of HLM half-life data, integration of broader metabolic systems, and improved uncertainty estimation to enhance model reliability across a wider chemical space.

## List of abbreviations

HLM: Human liver microsomes
RMSE: Root mean squared error
R^2^: Coefficient of determination
SHAP: Shapley Additive Explanations
QSAR: Quantitative structure–activity relationship
ML: Machine learning
KCB: Korea Chemical Bank
MLP: Multi-layer perceptron
k-NN: k-nearest neighbors
KRR: Kernel ridge regression
SVM: Support vector machine
RF: Random forest
CMPNN: Communicative message passing neural network
MSE: Mean squared error
MAE: Mean absolute error
AD: Applicability domain
SDC: Sum of distance-weighted contribution
PPV: Positive predictive value
NPV: Negative predictive value
MCC: Matthews correlation coefficient
ROC-AUC: Area under the receiver operating characteristic curve.

## Declarations

### Availability of data and materials

The source code, curated datasets, and trained model are freely available at https://github.com/joshua-416/PredHLM.

### Authors’ contributions

J.J. and K.S.O. designed the research; J.J. developed the prediction model; J.Y.R. contributed to model validation; J.J., N.C.C., and K.S.O. analyzed the data; all authors have read and approved the manuscript.

## Acknowledgments

Not applicable.

## References

1. Elaut G, Papeleu P, Vinken M, Henkens T, Snykers S, Vanhaecke T, Rogiers V (2006) Hepatocytes in suspension. In: Cytochrome P450 Protocols. Springer, pp 255–263.

2. Sakiyama Y, Yuki H, Moriya T, Hattori K, Suzuki M, Shimada K, Honma T (2008) Predicting human liver microsomal stability with machine learning techniques. Journal of molecular graphics and Modelling 26:907–915.

3. Schwaighofer A, Schroeter T, Mika S, Hansen K, Laak At, Lienau P, Reichel A, Heinrich N, Müller K-R (2008) A probabilistic approach to classifying metabolic stability. Journal of chemical information and modeling 48:785–796.

4. Hu Y, Unwalla R, Denny RA, Bikker J, Di L, Humblet C (2010) Development of QSAR models for microsomal stability: identification of good and bad structural features for rat, human and mouse microsomal stability. Journal of computer-aided molecular design 24:23–35.

5. Liu R, Schyman P, Wallqvist A (2015) Critically assessing the predictive power of QSAR models for human liver microsomal stability. Journal of Chemical Information and Modeling 55:1566–1575.

6. Venkatraman V (2021) FP-ADMET: a compendium of fingerprint-based ADMET prediction models. Journal of cheminformatics 13:75.

7. Ryu JY, Lee JH, Lee BH, Song JS, Ahn S, Oh K-S (2021) PredMS: a random Forest model for predicting metabolic stability of drug candidates in human liver microsomes. Bioinformatics 38:364–368.

8. Li L, Lu Z, Liu G, Tang Y, Li W (2022) In silico prediction of human and rat liver microsomal stability via machine learning methods. Chemical Research in Toxicology 35:1614–1624.

9. Long T-Z, Jiang D-J, Shi S-H, Deng Y-C, Wang W-X, Cao D-S (2024) Enhancing multi-species liver microsomal stability prediction through artificial intelligence. Journal of Chemical Information and Modeling 64:3222–3236.

10. Siramshetty VB, Shah P, Kerns E, Nguyen K, Yu KR, Kabir M, Williams J, Neyra J, Southall N, Nguyễn Ð-T, Xu X (2020) Retrospective assessment of rat liver microsomal stability at NCATS: data and QSAR models. Scientific reports 10:20713.

11. Shah P, Siramshetty VB, Mathé E, Xu X (2024) Developing robust human liver microsomal stability prediction models: Leveraging inter-species correlation with rat data. Pharmaceutics 16:1257.

12. Aliagas I, Gobbi A, Heffron T, Lee M-L, Ortwine DF, Zak M, Khojasteh SC (2015) A probabilistic method to report predictions from a human liver microsomes stability QSAR model: a practical tool for drug discovery. Journal of Computer-Aided Molecular Design 29:327–338.

13. Park JH, Han R, Jang J, Kim J, Paik J, Heo J, Lee Y (2025) MetaboGNN: predicting liver metabolic stability with graph neural networks and cross-species data. Journal of cheminformatics 17:140.

14. Ulenberg S, Belka M, Król M, Herold F, Hewelt-Belka W, Kot-Wasik A, Bączek T (2015) Prediction of overall in vitro microsomal stability of drug candidates based on molecular modeling and support vector machines. Case study of novel arylpiperazines derivatives. PLoS One 10:e0122772.

15. Zakharov AV, Peach ML, Sitzmann M, Filippov IV, McCartney HJ, Smith LH, Pugliese A, Nicklaus MC (2012) Computational tools and resources for metabolism-related property predictions. 2. Application to prediction of half-life time in human liver microsomes. Future medicinal chemistry 4:1933–1944.

16. Podlewska S, Kafel R (2018) MetStabOn—online platform for metabolic stability predictions. International journal of molecular sciences 19:1040.

17. Rudik A, Stolbov L, Lagunin A, Filimonov D, Poroikov V (2026) MetaStab Analyzer: Classification and Regression Models for Metabolic Stability Prediction. Molecular Informatics 45:e70018.

18. Lundberg SM, Lee S-I (2017) A unified approach to interpreting model predictions. Advances in neural information processing systems 30.

19. Rogers D, Hahn M (2010) Extended-connectivity fingerprints. Journal of chemical information and modeling 50:742–754.

20. Moriwaki H, Tian Y-S, Kawashita N, Takagi T (2018) Mordred: a molecular descriptor calculator. Journal of cheminformatics 10:4.

21. Brown RD, Martin YC (1997) The information content of 2D and 3D structural descriptors relevant to ligand-receptor binding. Journal of Chemical Information and Computer Sciences 37:1–9.

22. Murtagh F (1991) Multilayer perceptrons for classification and regression. Neurocomputing 2:183–197.

23. Peterson LE (2009) K-nearest neighbor. Scholarpedia 4:1883.

24. Vovk V (2013) Kernel ridge regression. In: Empirical inference: Festschrift in honor of vladimir n vapnik. Springer, pp 105–116.

25. Hearst MA, Dumais ST, Osuna E, Platt J, Scholkopf B (1998) Support vector machines. IEEE Intelligent Systems and their applications 13:18–28.

26. Breiman L (2001) Random forests. Machine learning 45:5–32.

27. Ke G, Meng Q, Finley T, Wang T, Chen W, Ma W, Ye Q, Liu T-Y (2017) Lightgbm: A highly efficient gradient boosting decision tree. Advances in neural information processing systems 30.

28. Chen T, Guestrin C (2016) Xgboost: A scalable tree boosting system. In: Proceedings of the 22nd acm sigkdd international conference on knowledge discovery and data mining, pp 785–794.

29. Prokhorenkova L, Gusev G, Vorobev A, Dorogush AV, Gulin A (2018) CatBoost: unbiased boosting with categorical features. Advances in neural information processing systems 31.

30. Song Y, Zheng S, Niu Z, Fu Z-H, Lu Y, Yang Y (2020) Communicative representation learning on attributed molecular graphs. In: 29th International Joint Conference on Artificial Intelligence and the 17th Pacific Rim International Conference on Artificial Intelligence (IJCAI-PRICAI2020). International Joint Conferences on Artificial Intelligence Organization.

31. Vaswani A, Shazeer N, Parmar N, Uszkoreit J, Jones L, Gomez AN, Kaiser Ł, Polosukhin I (2017) Attention is all you need. Advances in neural information processing systems 30.

32. Akiba T, Sano S, Yanase T, Ohta T, Koyama M (2019) Optuna: A next-generation hyperparameter optimization framework. In: Proceedings of the 25th ACM SIGKDD international conference on knowledge discovery & data mining, pp 2623–2631.

33. Wu Z, Ramsundar B, Feinberg EN, Gomes J, Geniesse C, Pappu AS, Leswing K, Pande V (2018) MoleculeNet: a benchmark for molecular machine learning. Chemical science 9:513–530.

34. Bemis GW, Murcko MA (1996) The properties of known drugs. 1. Molecular frameworks. Journal of medicinal chemistry 39:2887–2893.

35. Pedregosa F, Varoquaux G, Gramfort A, Michel V, Thirion B, Grisel O, Blondel M, Prettenhofer P, Weiss R, Dubourg V, et al (2011) Scikit-learn: Machine learning in Python. Journal of machine learning research 12:2825–2830.

36. Bajusz D, Rácz A, Héberger K (2015) Why is Tanimoto index an appropriate choice for fingerprint-based similarity calculations? Journal of cheminformatics 7:20.

37. Liu R, Wallqvist A (2018) Molecular similarity-based domain applicability metric efficiently identifies out-of-domain compounds. Journal of chemical information and modeling 59:181–189.

38. Lewis DF, Dickins M (2003) Baseline lipophilicity relationships in human cytochromes P450 associated with drug metabolism. Drug metabolism reviews 35:1– 18.

39. Waterhouse RN (2003) Determination of lipophilicity and its use as a predictor of blood–brain barrier penetration of molecular imaging agents. Molecular Imaging & Biology 5:376–389.

40. Zhang Z, Tang W (2018) Drug metabolism in drug discovery and development. Acta Pharmaceutica Sinica B 8:721–732.

41. Xu W, Wang W, Liu T, Xie J, Zhu C (2019) Late-stage trifluoromethylthiolation of benzylic CH bonds. Nature Communications 10:4867.

42. Brunelli F, Ceresa C, Fracchia L, Tron GC, Aprile S (2022) Expanding the chemical space of drug-like Passerini compounds: Can α-acyloxy carboxamides be considered hard drugs? ACS Medicinal Chemistry Letters 13:1898–1904.

43. Purser S, Moore PR, Swallow S, Gouverneur V (2008) Fluorine in medicinal chemistry. Chemical Society Reviews 37:320–330. doi:10.1039/B610213C.

44. Lassalas P, Gay B, Lasfargeas C, James MJ, Tran V, Vijayendran KG, Brunden KR, Kozlowski MC, Thomas CJ, Smith III AB, et al (2016) Structure property relationships of carboxylic acid isosteres. Journal of medicinal chemistry 59:3183–3203.

