## Supplementary Information for "PredHLM: quantitative and interpretable prediction of metabolic half-life in human liver microsomes"

**Supplementary Methods - Data curation and exclusion criteria**

Beyond the non-HLM systems described in the main text, ChEMBL-derived records were excluded if they represented any of the following assays or conditions outside phase I HLM parent-compound depletion:

- in vivo dosing experiments;
- enzyme-specific CYP binding or activity assays;
- prodrug activation;
- metabolite formation;
- clearance-only endpoints;
- monoamine oxidase (MAO) activity assays;
- UDPGA/glucuronidation (phase II) conditions;
- heat-inactivated microsomal controls.

Table S1. List of the retained RDKit2D and Mordred molecular descriptors used for model development

| Descriptor | Descriptor Name |
| --- | --- |
| RDKit2d (196) | MaxAbsEStateIndex, MaxEStateIndex, MinAbsEStateIndex, MinEStateIndex, qed, MolWt, HeavyAtomMolWt, ExactMolWt, NumValenceElectrons, NumRadicalElectrons, FpDensityMorgan1, FpDensityMorgan2, FpDensityMorgan3, AvgIpc, BalabanJ, BertzCT, Chi0, Chi0n, Chi0v, Chi1, Chi1n, Chi1v, Chi2n, Chi2v, Chi3n, Chi3v, Chi4n, Chi4v, HallKierAlpha, Kappa1, Kappa2, Kappa3, LabuteASA, PEOE_VSA1, PEOE_VSA10, PEOE_VSA11, PEOE_VSA12, PEOE_VSA13, PEOE_VSA14, PEOE_VSA2, PEOE_VSA3, PEOE_VSA4, PEOE_VSA5, PEOE_VSA6, PEOE_VSA7, PEOE_VSA8, PEOE_VSA9, SMR_VSA1, SMR_VSA10, SMR_VSA2, SMR_VSA3, SMR_VSA4, SMR_VSA5, SMR_VSA6, SMR_VSA7, SMR_VSA8, SMR_VSA9, SlogP_VSA1, SlogP_VSA10, SlogP_VSA11, SlogP_VSA12, SlogP_VSA2, SlogP_VSA3, SlogP_VSA4, SlogP_VSA5, SlogP_VSA6, SlogP_VSA7, SlogP_VSA8, SlogP_VSA9, TPSA, EState_VSA1, EState_VSA10, EState_VSA11, EState_VSA2, EState_VSA3, EState_VSA4, EState_VSA5, EState_VSA6, EState_VSA7, EState_VSA8, EState_VSA9, VSA_EState1, VSA_EState10, VSA_EState2, VSA_EState3, VSA_EState4, VSA_EState5, VSA_EState6, VSA_EState7, VSA_EState8, VSA_EState9, FractionCSP3, HeavyAtomCount, NHOHCount, NOCount, NumAliphaticCarbocycles, NumAliphaticHeterocycles, NumAliphaticRings, NumAromaticCarbocycles, NumAromaticHeterocycles, NumAromaticRings, NumHAcceptors, NumHDonors, NumHeteroatoms, NumRotatableBonds, NumSaturatedCarbocycles, NumSaturatedHeterocycles, NumSaturatedRings, RingCount, MolLogP, MolMR, fr_Al_COO, fr_Al_OH, fr_Al_OH_noTert, fr_ArN, fr_Ar_COO, fr_Ar_N, fr_Ar_NH, fr_Ar_OH, fr_COO, fr_COO2, fr_C_O, fr_C_O_noCOO, fr_C_S, fr_HOCCN, fr_Imine, fr_NH0, fr_NH1, fr_NH2, fr_N_O, fr_Ndealkylation1, fr_Ndealkylation2, fr_Nhpyrrole, fr_SH, fr_aldehyde, fr_alkyl_carbamate, fr_alkyl_halide, fr_allylic_oxid, fr_amide, fr_amidine, fr_aniline, fr_aryl_methyl, fr_azide, fr_azo, fr_barbitur, fr_benzene, fr_benzodiazepine, fr_bicyclic, fr_diazo, fr_dihydropyridine, fr_epoxide, fr_ester, fr_ether, fr_furan, fr_guanido, fr_halogen, fr_hdrzine, fr_hdrzone, fr_imidazole, fr_imide, fr_isocyan, fr_isothiocyan, fr_ketone, fr_ketone_Topliss, fr_lactam, fr_lactone, fr_methoxy, fr_morpholine, fr_nitrile, fr_nitro, fr_nitro_arom, fr_nitro_arom_nonortho, fr_nitroso, fr_oxazole, fr_oxime, fr_para_hydroxylation, fr_phenol, fr_phenol_noOrthoHbond, fr_phos_acid, fr_phos_ester, fr_piperdine, fr_piperzine, fr_priamide, fr_prisulfonamd, fr_pyridine, fr_quatN, fr_sulfide, fr_sulfonamd, fr_sulfone, fr_term_acetylene, fr_tetrazole, fr_thiazole, fr_thiocyan, fr_thiophene, fr_unbrch_alkane, fr_urea |
| Mordred (993) | ABC, ABCGG, nAcid, nBase, nAromAtom, nAromBond, nAtom, nHeavyAtom, nSpiro, nBridgehead, nHetero, nH, nB, nC, nN, nO, nS, nP, nF, nCl, nBr, nI, nX, ATS0dv, ATS1dv, ATS2dv, ATS3dv, ATS4dv, ATS5dv, ATS6dv, ATS7dv, ATS8dv, ATS0d, ATS1d, ATS2d, ATS3d, ATS4d, ATS5d, ATS6d, ATS7d, ATS8d, ATS0Z, ATS1Z, ATS2Z, ATS3Z, ATS4Z, ATS5Z, ATS6Z, ATS7Z, ATS8Z, ATS0m, ATS1m, ATS2m, ATS3m, ATS4m, ATS5m, ATS6m, ATS7m, ATS8m, ATS0v, ATS1v, ATS2v, ATS3v, ATS4v, ATS5v, ATS6v, ATS7v, ATS8v, ATS0se, ATS1se, ATS2se, ATS3se, ATS4se, ATS5se, ATS6se, ATS7se, ATS8se, ATS0pe, ATS1pe, ATS2pe, ATS3pe, ATS4pe, ATS5pe, ATS6pe, ATS7pe, ATS8pe, ATS0are, ATS1are, ATS2are, ATS3are, ATS4are, ATS5are, ATS6are, ATS7are, ATS8are, ATS0p, ATS1p, ATS2p, ATS3p, ATS4p, ATS5p, ATS6p, ATS7p, ATS8p, ATS0i, ATS1i, ATS2i, ATS3i, ATS4i, ATS5i, ATS6i, ATS7i, ATS8i, AATS0dv, AATS1dv, AATS2dv, AATS3dv, AATS4dv, AATS5dv, AATS0d, AATS1d, AATS2d, AATS3d, AATS4d, AATS5d, AATS0Z, AATS1Z, AATS2Z, AATS3Z, AATS4Z, AATS5Z, AATS0m, AATS1m, AATS2m, AATS3m, AATS4m, AATS5m, AATS0v, AATS1v, AATS2v, AATS3v, AATS4v, AATS5v, AATS0se, AATS1se, AATS2se, AATS3se, AATS4se, AATS5se, AATS0pe, AATS1pe, AATS2pe, AATS3pe, AATS4pe, AATS5pe, AATS0are, AATS1are, AATS2are, AATS3are, AATS4are, AATS5are, AATS0p, AATS1p, AATS2p, AATS3p, AATS4p, AATS5p, AATS0i, AATS1i, AATS2i, AATS3i, AATS4i, AATS5i, ATSC0dv, ATSC1dv, ATSC2dv, ATSC3dv, ATSC4dv, ATSC5dv, ATSC6dv, ATSC7dv, ATSC8dv, ATSC0d, ATSC1d, ATSC2d, ATSC3d, ATSC4d, ATSC5d, ATSC6d, ATSC7d, ATSC8d, ATSC0Z, ATSC1Z, ATSC2Z, ATSC3Z, ATSC4Z, ATSC5Z, ATSC6Z, ATSC7Z, ATSC8Z, ATSC0m, ATSC1m, ATSC2m, ATSC3m, ATSC4m, ATSC5m, ATSC6m, ATSC7m, ATSC8m, ATSC0v, ATSC1v, ATSC2v, ATSC3v, ATSC4v, ATSC5v, ATSC6v, ATSC7v, ATSC8v, ATSC0se, ATSC1se, ATSC2se, ATSC3se, ATSC4se, ATSC5se, ATSC6se, ATSC7se, ATSC8se, ATSC0pe, ATSC1pe, ATSC2pe, ATSC3pe, ATSC4pe, ATSC5pe, ATSC6pe, ATSC7pe, ATSC8pe, ATSC0are, ATSC1are, ATSC2are, ATSC3are, ATSC4are, ATSC5are, ATSC6are, ATSC7are, ATSC8are, ATSC0p, ATSC1p, ATSC2p, ATSC3p, ATSC4p, ATSC5p, ATSC6p, ATSC7p, ATSC8p, ATSC0i, ATSC1i, ATSC2i, ATSC3i, ATSC4i, ATSC5i, ATSC6i, ATSC7i, ATSC8i, AATSC0dv, AATSC1dv, AATSC2dv, AATSC3dv, AATSC4dv, AATSC5dv, AATSC0d, AATSC1d, AATSC2d, AATSC3d, AATSC4d, AATSC5d, AATSC0Z, AATSC1Z, AATSC2Z, AATSC3Z, AATSC4Z, AATSC5Z, AATSC0m, AATSC1m, AATSC2m, AATSC3m, AATSC4m, AATSC5m, AATSC0v, AATSC1v, AATSC2v, AATSC3v, AATSC4v, AATSC5v, AATSC0se, AATSC1se, AATSC2se, AATSC3se, AATSC4se, AATSC5se, AATSC0pe, AATSC1pe, AATSC2pe, AATSC3pe, AATSC4pe, AATSC5pe, AATSC0are, AATSC1are, AATSC2are, AATSC3are, AATSC4are, AATSC5are, AATSC0p, AATSC1p, AATSC2p, AATSC3p, AATSC4p, AATSC5p, AATSC0i, AATSC1i, AATSC2i, AATSC3i, AATSC4i, AATSC5i, MATS1dv, MATS2dv, MATS3dv, MATS4dv, MATS5dv, MATS1d, MATS2d, MATS3d, MATS4d, MATS5d, MATS1Z, MATS2Z, MATS3Z, MATS4Z, MATS5Z, MATS1m, MATS2m, MATS3m, MATS4m, MATS5m, MATS1v, MATS2v, MATS3v, MATS4v, MATS5v, MATS1se, MATS2se, MATS3se, MATS4se, MATS5se, MATS1pe, MATS2pe, MATS3pe, MATS4pe, MATS5pe, MATS1are, MATS2are, MATS3are, MATS4are, MATS5are, MATS1p, MATS2p, MATS3p, MATS4p, MATS5p, MATS1i, MATS2i, MATS3i, MATS4i, MATS5i, GATS1dv, GATS2dv, GATS3dv, GATS4dv, GATS5dv, GATS1d, GATS2d, GATS3d, GATS4d, GATS5d, GATS1Z, GATS2Z, GATS3Z, GATS4Z, GATS5Z, GATS1m, GATS2m, GATS3m, GATS4m, GATS5m, GATS1v, GATS2v, GATS3v, GATS4v, GATS5v, GATS1se, GATS2se, GATS3se, GATS4se, GATS5se, GATS1pe, GATS2pe, GATS3pe, GATS4pe, GATS5pe, GATS1are, GATS2are, GATS3are, GATS4are, GATS5are, GATS1p, GATS2p, GATS3p, GATS4p, GATS5p, GATS1i, GATS2i, GATS3i, GATS4i, GATS5i, BalabanJ, BertzCT, nBonds, nBondsO, nBondsS, nBondsD, nBondsT, nBondsA, nBondsM, nBondsKS, nBondsKD, C1SP1, C2SP1, C1SP2, C2SP2, C3SP2, C1SP3, C2SP3, C3SP3, C4SP3, HybRatio, FCSP3, Xch-3d, Xch-4d, Xch-5d, Xch-6d, Xch-7d, Xch-3dv, Xch-4dv, Xch-5dv, Xch-6dv, Xch-7dv, Xc-3d, Xc-4d, Xc-5d, Xc-6d, Xc-3dv, Xc-4dv, Xc-5dv, Xc-6dv, Xpc-4d, Xpc-5d, Xpc-6d, Xpc-4dv, Xpc-5dv, Xpc-6dv, Xp-1d, Xp-2d, Xp-3d, Xp-4d, Xp-5d, Xp-6d, Xp-7d, AXp-1d, AXp-2d, AXp-3d, AXp-4d, Xp-1dv, Xp-2dv, Xp-3dv, Xp-4dv, Xp-5dv, Xp-6dv, Xp-7dv, AXp-1dv, AXp-2dv, AXp-3dv, AXp-4dv, SZ, Sm, Sv, Sse, Spe, Sare, Sp, Si, MZ, Mm, Mv, Mse, Mpe, Mare, Mp, Mi, NsLi, NssBe, NssssBe, NssBH, NsssB, NssssB, NsCH3, NdCH2, NssCH2, NtCH, NdsCH, NaaCH, NsssCH, NddC, NtsC, NdssC, NaasC, NaaaC, NssssC, NsNH3, NsNH2, NssNH2, NdNH, NssNH, NaaNH, NtN, NsssNH, NdsN, NaaN, NsssN, NddsN, NaasN, NssssN, NsOH, NdO, NssO, NaaO, NsF, NsSiH3, NssSiH2, NsssSiH, NssssSi, NsPH2, NssPH, NsssP, NdsssP, NsssssP, NsSH, NdS, NssS, NaaS, NdssS, NddssS, NsCl, NsGeH3, NssGeH2, NsssGeH, NssssGe, NsAsH2, NssAsH, NsssAs, NsssdAs, NsssssAs, NsSeH, NdSe, NssSe, NaaSe, NdssSe, NddssSe, NsBr, NsSnH3, NssSnH2, NsssSnH, NssssSn, NsI, NsPbH3, NssPbH2, NsssPbH, NssssPb, SsLi, SssBe, SssssBe, SssBH, SsssB, SssssB, SsCH3, SdCH2, SssCH2, StCH, SdsCH, SaaCH, SsssCH, SddC, StsC, SdssC, SaasC, SaaaC, SssssC, SsNH3, SsNH2, SssNH2, SdNH, SssNH, SaaNH, StN, SsssNH, SdsN, SaaN, SsssN, SddsN, SaasN, SssssN, SsOH, SdO, SssO, SaaO, SsF, SsSiH3, SssSiH2, SsssSiH, SssssSi, SsPH2, SssPH, SsssP, SdsssP, SsssssP, SsSH, SdS, SssS, SaaS, SdssS, SddssS, SsCl, SsGeH3, SssGeH2, SsssGeH, SssssGe, SsAsH2, SssAsH, SsssAs, SsssdAs, SsssssAs, SsSeH, SdSe, SssSe, SaaSe, SdssSe, SddssSe, SsBr, SsSnH3, SssSnH2, SsssSnH, SssssSn, SsI, SsPbH3, SssPbH2, SsssPbH, SssssPb, ECIndex, fragCpx, fMF, nHBAcc, nHBDon, IC0, IC1, IC2, IC3, IC4, IC5, TIC0, TIC1, TIC2, TIC3, TIC4, TIC5, SIC0, SIC1, SIC2, SIC3, SIC4, SIC5, BIC0, BIC1, BIC2, BIC3, BIC4, BIC5, CIC0, CIC1, CIC2, CIC3, CIC4, CIC5, MIC0, MIC1, MIC2, MIC3, MIC4, MIC5, ZMIC0, ZMIC1, ZMIC2, ZMIC3, ZMIC4, ZMIC5, Kier1, Kier2, Kier3, Lipinski, GhoseFilter, FilterItLogS, VMcGowan, LabuteASA, PEOE_VSA1, PEOE_VSA2, PEOE_VSA3, PEOE_VSA4, PEOE_VSA5, PEOE_VSA6, PEOE_VSA7, PEOE_VSA8, PEOE_VSA9, PEOE_VSA10, PEOE_VSA11, PEOE_VSA12, PEOE_VSA13, SMR_VSA1, SMR_VSA2, SMR_VSA3, SMR_VSA4, SMR_VSA5, SMR_VSA6, SMR_VSA7, SMR_VSA8, SMR_VSA9, SlogP_VSA1, SlogP_VSA2, SlogP_VSA3, SlogP_VSA4, SlogP_VSA5, SlogP_VSA6, SlogP_VSA7, SlogP_VSA8, SlogP_VSA9, SlogP_VSA10, SlogP_VSA11, EState_VSA1, EState_VSA2, EState_VSA3, EState_VSA4, EState_VSA5, EState_VSA6, EState_VSA7, EState_VSA8, EState_VSA9, EState_VSA10, VSA_EState1, VSA_EState2, VSA_EState3, VSA_EState4, VSA_EState5, VSA_EState6, VSA_EState7, VSA_EState8, VSA_EState9, MPC2, MPC3, MPC4, MPC5, MPC6, MPC7, MPC8, MPC9, MPC10, TMPC10, piPC1, piPC2, piPC3, piPC4, piPC5, piPC6, piPC7, piPC8, piPC9, piPC10, TpiPC10, apol, bpol, nRing, n3Ring, n4Ring, n5Ring, n6Ring, n7Ring, n8Ring, n9Ring, n10Ring, n11Ring, n12Ring, nG12Ring, nHRing, n3HRing, n4HRing, n5HRing, n6HRing, n7HRing, n8HRing, n9HRing, n10HRing, n11HRing, n12HRing, nG12HRing, naRing, n3aRing, n4aRing, n5aRing, n6aRing, n7aRing, n8aRing, n9aRing, n10aRing, n11aRing, n12aRing, nG12aRing, naHRing, n3aHRing, n4aHRing, n5aHRing, n6aHRing, n7aHRing, n8aHRing, n9aHRing, n10aHRing, n11aHRing, n12aHRing, nG12aHRing, nARing, n3ARing, n4ARing, n5ARing, n6ARing, n7ARing, n8ARing, n9ARing, n10ARing, n11ARing, n12ARing, nG12ARing, nAHRing, n3AHRing, n4AHRing, n5AHRing, n6AHRing, n7AHRing, n8AHRing, n9AHRing, n10AHRing, n11AHRing, n12AHRing, nG12AHRing, nFRing, n4FRing, n5FRing, n6FRing, n7FRing, n8FRing, n9FRing, n10FRing, n11FRing, n12FRing, nG12FRing, nFHRing, n4FHRing, n5FHRing, n6FHRing, n7FHRing, n8FHRing, n9FHRing, n10FHRing, n11FHRing, n12FHRing, nG12FHRing, nFaRing, n4FaRing, n5FaRing, n6FaRing, n7FaRing, n8FaRing, n9FaRing, n10FaRing, n11FaRing, n12FaRing, nG12FaRing, nFaHRing, n4FaHRing, n5FaHRing, n6FaHRing, n7FaHRing, n8FaHRing, n9FaHRing, n10FaHRing, n11FaHRing, n12FaHRing, nG12FaHRing, nFARing, n4FARing, n5FARing, n6FARing, n7FARing, n8FARing, n9FARing, n10FARing, n11FARing, n12FARing, nG12FARing, nFAHRing, n4FAHRing, n5FAHRing, n6FAHRing, n7FAHRing, n8FAHRing, n9FAHRing, n10FAHRing, n11FAHRing, n12FAHRing, nG12FAHRing, nRot, RotRatio, SLogP, SMR, TopoPSA(NO), TopoPSA, GGI1, GGI2, GGI3, GGI4, GGI5, GGI6, GGI7, GGI8, GGI9, GGI10, JGI1, JGI2, JGI3, JGI4, JGI5, JGI6, JGI7, JGI8, JGI9, JGI10, JGT10, Diameter, Radius, TopoShapeIndex, PetitjeanIndex, VAdjMat, MWC01, MWC02, MWC03, MWC04, MWC05, MWC06, MWC07, MWC08, MWC09, MWC10, TMWC10, SRW02, SRW03, SRW04, SRW05, SRW06, SRW07, SRW08, SRW09, SRW10, TSRW10, MW, AMW, WPath, WPol, Zagreb1, Zagreb2, mZagreb2 |

Table S2. Hyperparameter search spaces for the machine-learning models. All hyperparameters were optimized using the Optuna framework with the Tree-structured Parzen Estimator (TPE) sampler. “Type” indicates the sampling distribution: integers and floats were drawn uniformly, while “log” denotes log-uniform sampling. Categorical parameters were drawn from the listed discrete sets. For the gradient-boosted decision tree models (CatBoost, LightGBM, XGBoost) the number of estimators was fixed at 2000 and the effective number of trees was determined by early stopping (50-round patience on the validation set), so it is not listed as a tuned hyperparameter.

| Model | Hyperparameter | Search space | Type |
| --- | --- | --- | --- |
| CatBoost | *depth* | 3 - 12 | integer |
|  | *learning_rate* | 0.005 - 0.15 | float, log |
|  | *l2_leaf_reg* | 1.0 - 50.0 | float, log |
|  | *min_child_samples* | 10 - 150 | integer |
|  | *colsample_bylevel* | 0.5 - 1.0 | float ᵃ |
| LightGBM | *num_leaves* | 15 - 63 | integer |
|  | *max_depth* | 3 - 12 | integer |
|  | *learning_rate* | 0.005 - 0.2 | float, log |
|  | *min_child_samples* | 10 - 150 | integer |
|  | *reg_alpha* | 1×10^-4^ - 10.0 | float, log |
|  | *reg_lambda* | 1×10^-4^- 10.0 | float, log |
|  | *subsample* | 0.5 - 1.0 | float |
|  | *colsample_bytree* | 0.5 - 1.0 | float |
| XGBoost | *max_depth* | 3 - 12 | integer |
|  | *learning_rate* | 0.005 - 0.2 | float, log |
|  | *min_child_weight* | 1 - 50 | integer |
|  | *subsample* | 0.5 - 1.0 | float |
|  | *colsample_bytree* | 0.5 - 1.0 | float |
|  | *reg_alpha* | 1×10^-4^ - 10.0 | float, log |
|  | *reg_lambda* | 1×10^-4^ - 10.0 | float, log |
|  | *gamma* | 0.0 - 0.5 | float |
| Random Forest | *n_estimators* | 200 - 1500 | integer |
|  | *max_depth* | 5 - 30 | integer |
|  | *min_samples_leaf* | 2 - 30 | integer |
|  | *min_samples_split* | 2 - 30 | integer |
|  | *max_features* | {sqrt, 0.3, 0.5} | categorical |
| k-NN | *n_neighbors* | 3 - 25 | integer |
|  | *weights* | {uniform, distance} | categorical |
|  | *metric* | {euclidean, manhattan} | categorical |
| SVM | *C* | 0.01 - 100 | float, log |
|  | *gamma* | {scale, auto} | categorical |
|  | *kernel* | {rbf, linear} | categorical |
|  | *epsilon* | 0.01 - 1.0 | float, log |
| KRR | *alpha* | 0.01 - 100 | float, log |
|  | *kernel* | {linear, rbf, polynomial} | categorical |
|  | *gamma* | 1×10^-4^ - 1.0 | float, log ᵇ |
| MLP | *n_layers* | 1 - 3 | integer |
|  | *layer_size* | 64 - 512 | integer |
|  | *activation* | {relu, tanh} | categorical |
|  | *alpha* | 1×10^-5^ - 0.1 | float, log |
|  | *learning_rate_init* | 1×10^-4^ - 1×10⁻² | float, log |
| CMPNN | *hidden_size* | {128, 256, 300, 384, 512} | categorical |
|  | *depth* | 2 - 6 | integer |
|  | *dropout* | 0.0 - 0.5 | float |
|  | *ffn_num_layers* | 1 - 3 | integer |
|  | *ffn_hidden_size* | {128, 256, 300, 384, 512} | categorical |
|  | *activation* | {ReLU, LeakyReLU, PReLU, ELU} | categorical |
|  | *batch_size* | {32, 64, 128, 256} | categorical |
|  | *max_lr* | 1×10^-4^ - 1×10⁻² | float, log |
|  | *init_lr_ratio* | 0.01 - 0.5 | float, log |
|  | *final_lr_ratio* | 0.001 - 0.5 | float, log |
|  | *warmup_epochs* | {1.0, 2.0, 3.0} | categorical |
|  | *attention_dim^a^* | {32, 64, 128, 256} | categorical |

^a^ Tuned only for models with the self-attention readout

Table S3. Optimal hyperparameters selected by Optuna (TPE sampler, 5-fold cross-validation) for each model and molecular feature set. Each value is the best configuration identified on the cross-validation objective

| Model | Hyperparameter | RDKit2D | Mordred | ECFP4 | MACCS |
| --- | --- | --- | --- | --- | --- |
| CatBoost | *depth* | 11 | 8 | 12 | 10 |
|  | *learning_rate* | 0.02524 | 0.04092 | 0.042 | 0.02004 |
|  | *l2_leaf_reg* | 4.081 | 1.904 | 3.934 | 1.454 |
|  | *min_child_samples* | 116 | 37 | 137 | 65 |
|  | *colsample_bylevel* | 0.7698 | 0.5449 | 0.6856 | 0.7707 |
| LightGBM | *num_leaves* | 61 | 59 | 62 | 58 |
|  | *max_depth* | 11 | 9 | 10 | 12 |
|  | *learning_rate* | 0.01333 | 0.02087 | 0.02566 | 0.04886 |
|  | *min_child_samples* | 30 | 10 | 16 | 11 |
|  | *reg_alpha* | 0.3663 | 2.144 | 1.684 | 0.001619 |
|  | *reg_lambda* | 1.583 | 0.005555 | 0.8655 | 0.2827 |
|  | *subsample* | 0.8802 | 0.5606 | 0.807 | 0.8684 |
|  | *colsample_bytree* | 0.7152 | 0.5632 | 0.6227 | 0.5287 |
| XGBoost | *max_depth* | 12 | 10 | 10 | 12 |
|  | *learning_rate* | 0.01954 | 0.01298 | 0.01326 | 0.006356 |
|  | *min_child_weight* | 20 | 33 | 8 | 2 |
|  | *subsample* | 0.7836 | 0.7311 | 0.9121 | 0.6466 |
|  | *colsample_bytree* | 0.7428 | 0.8582 | 0.5813 | 0.5949 |
|  | *reg_alpha* | 1.135 | 0.01273 | 0.0177 | 0.05979 |
|  | *reg_lambda* | 0.05496 | 4.18 | 0.0132 | 0.000344 |
|  | *gamma* | 0.07772 | 0.07583 | 0.01601 | 0.0246 |
| Random Forest | *n_estimators* | 1287 | 588 | 832 | 490 |
|  | *max_depth* | 25 | 30 | 30 | 27 |
|  | *min_samples_leaf* | 2 | 2 | 2 | 2 |
|  | *min_samples_split* | 6 | 5 | 9 | 4 |
|  | *max_features* | 0.5 | 0.5 | 0.5 | 0.3 |
| k-NN | *n_neighbors* | 14 | 12 | 9 | 15 |
|  | *weights* | distance | distance | distance | distance |
|  | *metric* | manhattan | manhattan | manhattan | manhattan |
| SVM | *C* | 1.842 | 2.308 | 1.232 | 1.342 |
|  | *gamma* | scale | scale | scale | scale |
|  | *kernel* | rbf | rbf | rbf | rbf |
|  | *epsilon* | 0.1588 | 0.1374 | 0.1061 | 0.1419 |
| KRR | *alpha* | 0.1276 | 0.1039 | 11.44 | 66.59 |
|  | *kernel* | rbf | rbf | polynomial | polynomial |
|  | *gamma* | 0.00264 | 0.000465 | 0.001446 | 0.02694 |
| MLP | *n_layers* | 2 | 3 | 3 | 2 |
|  | *layer_size* | 403 | 496 | 499 | 511 |
|  | *activation* | relu | relu | relu | relu |
|  | *alpha* | 0.000205 | 0.08996 | 0.001649 | 0.04583 |
|  | *learning_rate_init* | 0.000134 | 0.000373 | 0.000348 | 0.000547 |

Table S4. Optimal hyperparameters selected by Optuna (TPE sampler, 5-fold cross-validation) for CMPNN model and its variants. Each value is the best configuration identified on the cross-validation objective

| Model | Hyperparameter | value |
| --- | --- | --- |
| baseline | *hidden_size* | 384 |
|  | *depth* | 5 |
|  | *dropout* | 0.008294 |
|  | *ffn_num_layers* | 2 |
|  | *ffn_hidden_size* | 384 |
|  | *activation* | ReLU |
|  | *batch_size* | 32 |
|  | *max_lr* | 0.000236 |
|  | *init_lr_ratio* | 0.01173 |
|  | *final_lr_ratio* | 0.03934 |
|  | *warmup_epochs* | 1 |
|  | *attention_dim* | N/A |
| attention | *hidden_size* | 256 |
|  | *depth* | 3 |
|  | *dropout* | 0.08789 |
|  | *ffn_num_layers* | 1 |
|  | *ffn_hidden_size* | 384 |
|  | *activation* | PReLU |
|  | *batch_size* | 32 |
|  | *max_lr* | 0.000166 |
|  | *init_lr_ratio* | 0.3593 |
|  | *final_lr_ratio* | 0.3551 |
|  | *warmup_epochs* | 2 |
|  | *attention_dim* | 32 |
| baseline+rdkit2d | *hidden_size* | 384 |
|  | *depth* | 5 |
|  | *dropout* | 0.0584 |
|  | *ffn_num_layers* | 3 |
|  | *ffn_hidden_size* | 256 |
|  | *activation* | LeakyReLU |
|  | *batch_size* | 64 |
|  | *max_lr* | 0.000276 |
|  | *init_lr_ratio* | 0.06277 |
|  | *final_lr_ratio* | 0.01105 |
|  | *warmup_epochs* | 2 |
|  | *attention_dim* | N/A |
| attention+rdkit2d | *hidden_size* | 384 |
|  | *depth* | 2 |
|  | *dropout* | 0.2499 |
|  | *ffn_num_layers* | 3 |
|  | *ffn_hidden_size* | 512 |
|  | *activation* | ReLU |
|  | *batch_size* | 128 |
|  | *max_lr* | 0.001539 |
|  | *init_lr_ratio* | 0.1545 |
|  | *final_lr_ratio* | 0.004102 |
|  | *warmup_epochs* | 3 |
|  | *attention_dim* | 32 |
| baseline+mordred | *hidden_size* | 256 |
|  | *depth* | 2 |
|  | *dropout* | 0.2077 |
|  | *ffn_num_layers* | 2 |
|  | *ffn_hidden_size* | 256 |
|  | *activation* | ELU |
|  | *batch_size* | 64 |
|  | *max_lr* | 0.001501 |
|  | *init_lr_ratio* | 0.04061 |
|  | *final_lr_ratio* | 0.003684 |
|  | *warmup_epochs* | 1 |
|  | *attention_dim* | N/A |
| attention+mordred | *hidden_size* | 384 |
|  | *depth* | 3 |
|  | *dropout* | 0.2081 |
|  | *ffn_num_layers* | 3 |
|  | *ffn_hidden_size* | 256 |
|  | *activation* | LeakyReLU |
|  | *batch_size* | 64 |
|  | *max_lr* | 0.002274 |
|  | *init_lr_ratio* | 0.1056 |
|  | *final_lr_ratio* | 0.01699 |
|  | *warmup_epochs* | 3 |
|  | *attention_dim* | 128 |
| baseline+ECFP4 | *hidden_size* | 300 |
|  | *depth* | 2 |
|  | *dropout* | 0.2697 |
|  | *ffn_num_layers* | 2 |
|  | *ffn_hidden_size* | 512 |
|  | *activation* | ReLU |
|  | *batch_size* | 256 |
|  | *max_lr* | 0.001658 |
|  | *init_lr_ratio* | 0.3196 |
|  | *final_lr_ratio* | 0.003557 |
|  | *warmup_epochs* | 3 |
|  | *attention_dim* | N/A |
| attention+ECFP4 | *hidden_size* | 512 |
|  | *depth* | 2 |
|  | *dropout* | 0.2344 |
|  | *ffn_num_layers* | 2 |
|  | *ffn_hidden_size* | 256 |
|  | *activation* | PReLU |
|  | *batch_size* | 32 |
|  | *max_lr* | 0.000131 |
|  | *init_lr_ratio* | 0.2003 |
|  | *final_lr_ratio* | 0.1444 |
|  | *warmup_epochs* | 1 |
|  | *attention_dim* | 256 |
| baseline+maccs | *hidden_size* | 384 |
|  | *depth* | 3 |
|  | *dropout* | 0.06534 |
|  | *ffn_num_layers* | 3 |
|  | *ffn_hidden_size* | 384 |
|  | *activation* | LeakyReLU |
|  | *batch_size* | 64 |
|  | *max_lr* | 0.000164 |
|  | *init_lr_ratio* | 0.3107 |
|  | *final_lr_ratio* | 0.1452 |
|  | *warmup_epochs* | 1 |
|  | *attention_dim* | N/A |
| attention+maccs | *hidden_size* | 512 |
|  | *depth* | 5 |
|  | *dropout* | 0.09152 |
|  | *ffn_num_layers* | 2 |
|  | *ffn_hidden_size* | 300 |
|  | *activation* | LeakyReLU |
|  | *batch_size* | 32 |
|  | *max_lr* | 0.000404 |
|  | *init_lr_ratio* | 0.385 |
|  | *final_lr_ratio* | 0.001479 |
|  | *warmup_epochs* | 1 |
|  | *attention_dim* | 128 |

Table S5. Cross-validated regression performance of the three featurization schemes other than RDKit 2D (ECFP4 fingerprints, Mordred descriptors, and MACCS keys) evaluated for HLM half-life prediction. Values are reported as the mean ± standard deviation of RMSE, MAE, and R^2^ across 5-fold cross-validation; for each featurization scheme, the best-performing model is shown in bold

| Feature | Algorithm | RMSE (↓) | MAE (↓) | R^2^(↑) |
| --- | --- | --- | --- | --- |
| Mordred | MLP | 0.564 ± 0.013 | 0.421 ± 0.007 | 0.293 ± 0.030 |
|  | k-NN | 0.562 ± 0.005 | 0.424 ± 0.006 | 0.297 ± 0.010 |
|  | KRR | 0.537 ± 0.006 | 0.406 ± 0.005 | 0.358 ± 0.009 |
|  | SVM | 0.527 ± 0.006 | 0.392 ± 0.006 | 0.381 ± 0.009 |
|  | RF | 0.535 ± 0.005 | 0.407 ± 0.007 | 0.363 ± 0.008 |
|  | LightGBM | 0.515 ± 0.003 | 0.384 ± 0.005 | 0.411 ± 0.008 |
|  | CatBoost | 0.516 ± 0.004 | 0.387 ± 0.006 | 0.407 ± 0.005 |
|  | **XGBoost** | **0.514 ± 0.003** | **0.383 ± 0.005** | **0.412 ± 0.009** |
|  | CMPNN + feature | 0.539 ± 0.005 | 0.407 ± 0.004 | 0.353 ± 0.009 |
|  | CMPNN + attention + feature | 0.528 ± 0.003 | 0.396 ± 0.002 | 0.379 ± 0.008 |
| ECFP4 | MLP | 0.555 ± 0.009 | 0.414 ± 0.007 | 0.316 ± 0.017 |
|  | k-NN | 0.565 ± 0.007 | 0.425 ± 0.006 | 0.290 ± 0.006 |
|  | KRR | 0.527 ± 0.007 | 0.395 ± 0.007 | 0.381 ± 0.014 |
|  | SVM | 0.527 ± 0.007 | 0.392 ± 0.006 | 0.381 ± 0.013 |
|  | RF | 0.557 ± 0.008 | 0.427 ± 0.008 | 0.310 ± 0.011 |
|  | LightGBM | 0.535 ± 0.009 | 0.404 ± 0.008 | 0.364 ± 0.018 |
|  | CatBoost | 0.528 ± 0.007 | 0.397 ± 0.006 | 0.379 ± 0.014 |
|  | XGBoost | 0.534 ± 0.008 | 0.402 ± 0.008 | 0.366 ± 0.014 |
|  | CMPNN + feature | 0.526 ± 0.003 | 0.392 ± 0.004 | 0.385 ± 0.012 |
|  | **CMPNN + attention + feature** | **0.518 ± 0.004** | **0.385 ± 0.005** | **0.403 ± 0.015** |
| MACCS | MLP | 0.576 ± 0.007 | 0.441 ± 0.004 | 0.262 ± 0.006 |
|  | k-NN | 0.577 ± 0.006 | 0.437 ± 0.007 | 0.259 ± 0.013 |
|  | KRR | 0.548 ± 0.005 | 0.414 ± 0.005 | 0.331 ± 0.009 |
|  | SVM | 0.558 ± 0.005 | 0.422 ± 0.005 | 0.307 ± 0.005 |
|  | RF | 0.544 ± 0.007 | 0.412 ± 0.008 | 0.341 ± 0.009 |
|  | LightGBM | 0.545 ± 0.006 | 0.411 ± 0.006 | 0.339 ± 0.010 |
|  | CatBoost | 0.540 ± 0.005 | 0.408 ± 0.006 | 0.351 ± 0.011 |
|  | XGBoost | 0.539 ± 0.006 | 0.405 ± 0.007 | 0.354 ± 0.010 |
|  | **CMPNN + feature** | **0.529 ± 0.002** | **0.399 ± 0.002** | **0.378 ± 0.012** |
|  | CMPNN + attention + feature | 0.531 ± 0.004 | 0.401 ± 0.007 | 0.372 ± 0.014 |

Table S6. Regression performance of the selected RDKit 2D-XGBoost model under random and scaffold-based data splits. Cross-validation values are reported as mean ± standard deviation across the 5 folds. Independent-test values were obtained from the ensemble of the 5fold-specific models (averaged predictions) evaluated once on the test set, and are therefore reported as single values

| Split | Evaluation set | RMSE (↓) | MAE (↓) | R^2^(↑) |
| --- | --- | --- | --- | --- |
| Random split | Cross-validation | 0.515 ± 0.007 | 0.387 ± 0.007 | 0.410 ± 0.011 |
|  | Independent test set | 0.507 | 0.379 | 0.431 |
| Scaffold split | Cross-validation | 0.545 ± 0.005 | 0.419 ± 0.003 | 0.338 ± 0.017 |
|  | Independent test set | 0.536 | 0.412 | 0.369 |
